# Subcellularly Resolved 3D Translatome in Mouse Oocytes and Early Embryos

**DOI:** 10.64898/2026.07.31.742119

**Authors:** Jingyi Ren, Seth Furniss, Chengjie Zhou, Haowen Zhou, Yota Hagihara, Xiao Wang, Yi Zhang

## Abstract

Spatial patterning of mRNA translation is a fundamental process in early embryogenesis. Existing RNA translation profiling methods lack subcellular spatial resolution at the single-molecule level, limiting our understanding of spatial RNA biology in embryogenesis. To address this, we profiled the spatial translatome of intact mouse embryos at near-genomic scale by adapting RIBOmap and incorporating multiplexed organelle staining. In oocytes, 2-cell and 4-cell embryos, we systematically analyzed RNA translation across three spatial scales: organelle, subcellular, and intercellular. We found that functionally related genes exhibit spatially and temporally controlled translation patterns near distinct organelles. Using Harmonics, a graph signal processing framework, we demonstrate that embryo asymmetry emerges at the first cell division and is amplified at later stages. This work paves the way for comprehensively investigating the fundamental spatial post-transcriptional regulation at the earliest moments of mammalian life.

## Main Text

Visualizing mRNA transcripts across their translational states at single-molecule resolution in situ can reveal how post-transcriptional regulation operates spatially within cells (*1–3*). This approach is particularly informative for understanding dynamic translational remodeling at the beginning of mammalian life, during oocyte maturation and early embryonic development (*4*). For example, localization of maternal mRNA molecules in frog (*Xenopus*) (*5–7*) and fruit fly (*Drosophila melanogaster*) (*8*) oocytes has been shown to establish the foundation for early embryonic patterning. Meanwhile, studies in mammals have shown that maternal mRNA translation remains active from fully matured oocytes to early zygotic stages, even as the major transcription of new mRNAs has not yet been initiated. In mice, active translation of maternal mRNAs is essential for the maternal-to-zygotic transition, impacting processes like chromatin reprogramming and zygotic genome activation (ZGA) (*9–11*), while differential ribosome engagement of zygotic mRNAs exhibits allele-biased gene expression (*12*). However, these observations have been limited to bulk or low-input single-cell analyses that require cell lysis and dissociation of RNAs from embryos, inherently sacrificing subcellular spatial context about when and where mRNA translation occurs in individual intact samples (*9*, *10*, *12*). There is, therefore, a critical need for introducing new in situ technologies for translatome-wide profiling in a spatially and temporally resolved manner within intact oocytes and early embryos.

Imaging-based spatial omics technologies have enabled in situ sequencing of mRNAs in intact biological samples, preserving native subcellular architecture across complex tissues (*13*, *14*). Building on this, RIBOmap (*15*) extends spatial profiling to the translatome by selectively amplifying and sequencing ribosome-bound transcripts, enabling spatial mapping of hundreds to thousands of translating mRNAs. However, extending spatial translatomics to early developing embryos remains a significant technical challenge due to their fragility and spherical, whole-mount nature. Although spatial protocols such as RIBOmap have been successfully applied to cultured cells and solid tissue sections (*15–16*), adapting these to systems of oocytes and embryos presents several unique experimental hurdles that need to overcome to ensure robust RNA detection while enabling concurrent multiplexed organelle staining.

To address this gap, we have developed optimized versions of RIBOmap tailored for intact preimplantation embryos, enabling robust detection of ribosome-bound mRNAs in samples up to 80 *µ*m in z depth with multiplexed organelle staining to provide precise subcellular context. Using these experimental procedures, we systematically mapped the spatiotemporal landscape of the translatome in different developmental stages, revealing a coherent picture of how subcellular RNA localization is functionally organized. Building on this organelle-proximal organization, we further zoomed out to the cellular length scale and uncovered a dynamic spatial polarization of RNA patterns along the embryo’s geometry. We also developed Harmonics, a graph signal processing (*17*)-based framework, which enabled us to identify interior-shell RNA patterning in MII oocytes and asymmetric RNA translatome distributions within individual blastomeres. We further validated asymmetric localization of a transcription factor *Jdp2*, which shows that it biases cell fate toward the inner cell mass (ICM) lineage upon overexpression in one blastomere of the 2-cell embryo. Collectively, these findings reveal a multilayered spatial organization of RNA translation, from organelle-level compartmentalization to embryo-wide polarity and blastomere asymmetry, that may shape early cell fate decisions at molecular, subcellular, and intercellular levels.

### Adaptation of RIBOmap for intact pre-implantation embryos

Direct application of our previously established technique RIBOmap (*15*), optimized for cultured cells and tissues, to embryos poses several unique challenges. Here, we have optimized the technology in several ways: First, embryos are naturally non-adherent and do not firmly attach to glass surfaces, therefore failing to satisfy the solid-phase attachment prerequisite of the standard protocol. To overcome this, we directly embedded embryos into a thin layer of agarose gel prior to RIBOmap chemical processing **(Fig. 1A, Step A)**. Second, for high subcellular resolution mapping of RNA transcripts, a concurrent visualization of subcellular organelles will provide spatial context for transcript localization. Also, at the multicellular embryo stages, it is necessary to obtain spatial information of nuclei and cell membrane to enable cell segmentation. Here, we integrated the current procedure with simultaneous four-plex cellular organelle staining to expand the imaging modality of RIBOmap in embryos **(Fig. 1A, Step B)**. Third, the physical dimensions of oocytes and embryos (60-80 *μ*m in diameter) require a compatible protocol for 28S ribosome-bound RNA targeting using three-probe strategy, probe amplification and anchoring in 3D thick samples. Inspired by Deep-STARmap and Deep-RIBOmap (*16*), we optimized the protocol for DNA amplicon library preparation and volumetric imaging in 3D samples while preserving the capability to detect organelle localization **(Fig. 1A, Step C-D)**. Last, we read out the gene identity of each amplicon through barcode-based in situ sequencing as described previously **(Fig. 1A, Step E)** (*18*). Collectively, the implementation of these unique strategies allowed us to modify RIBOmap for 3D spatial translatome profiling in non-dissociated, intact embryos as a whole.

**Fig. 1.**
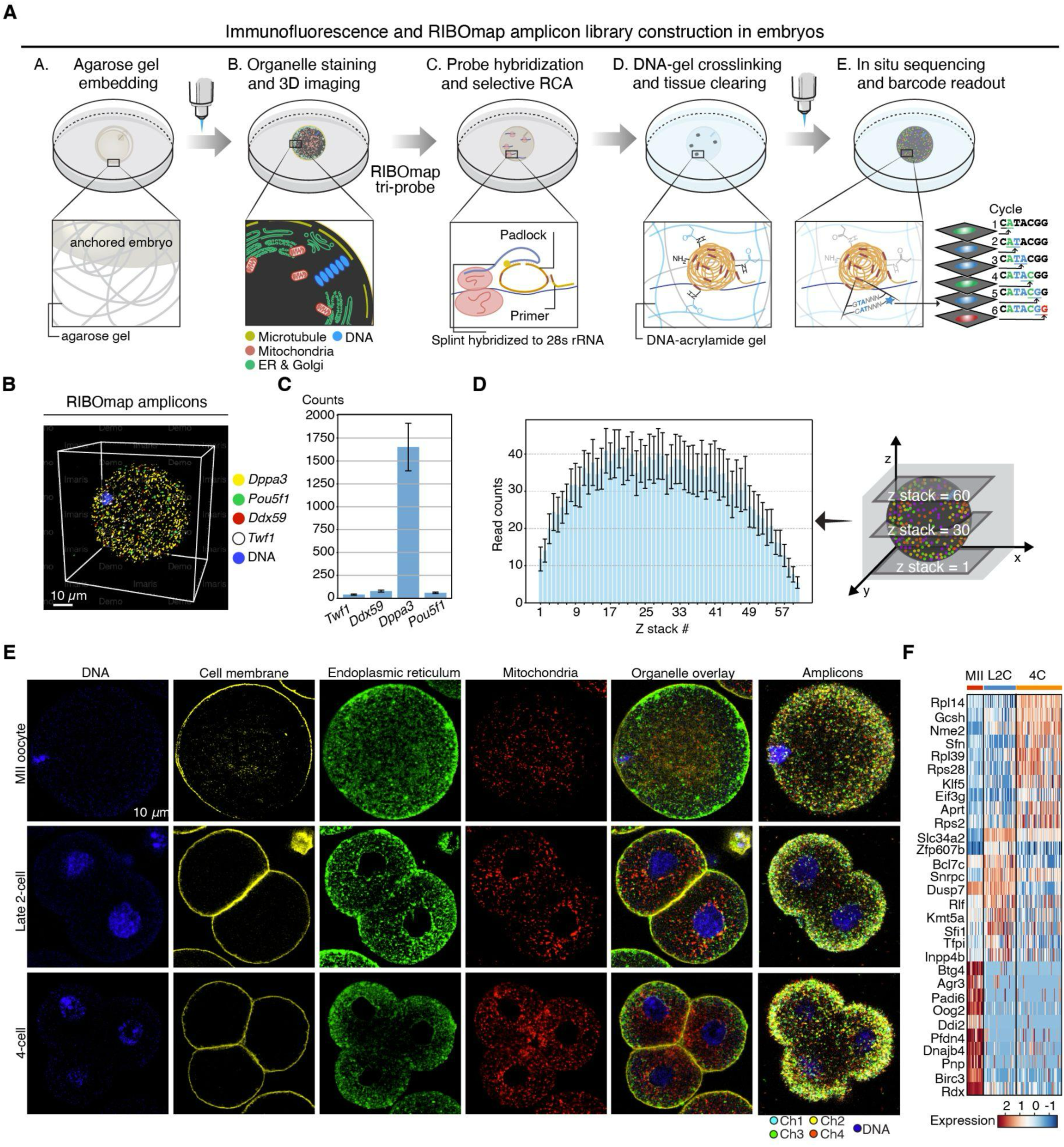
Optimized 3D RIBOmap enables multiplexed RNA translation mapping in oocytes and embryos. (**A**) Schematic overview of the method: intact embryos are freshly collected and anchored, followed by organelle staining and 3D imaging. Samples are then subjected to RIBOmap probe hybridization and library preparation, covalently embedded within a hydrogel matrix, optically cleared, and sequenced in situ(*14*). Detailed library preparation workflows are described in the Methods section. (**B**) 3D rendering of fluorescent oocyte images from 4-gene RIBOmap amplicons targeting *Twf1*, *Ddx59*, *Dppa3* and *Pou5f1*. (**C**) Averaged RIBOmap read counts per embryo for each gene across six samples. Data are presented as mean values ± s.d. (**D**) Read counts of four genes in (**b**) across the z-depth of embryos (left) and the illustration of z-stack imaging is shown (right). (**E**) Representative fluorescent embryo images of different stages of 4-plexed organelle detection to visualize nuclei, cortical actin filaments, endoplasmic reticulum, and mitochondria simultaneously, as well as RIBOmap sequencing (four color channels + DNA), all shown as one z-stack. Scale bar: 10 *μ*m. (**F**) Heatmap of normalized representative gene expression across developmental stages.

As a technical validation, we curated a list of four genes (*Dppa3, Pou5f1, Twf1,* and *Ddx59*) that show differential gene expression in the oocyte Metaphase II (MII) stage (**Fig. 1B**) (*9*, *10*).We simultaneously targeted the four genes with variable abundance (**Fig. 1C**), and demonstrated that the optimized methods enable robust and consistent signal detection across the full depth of z-stacks spanning 60-80 *µ*m embryo samples **(Fig. 1D)**. These results indicate that we have developed an optimized RIBOmap protocol that achieves high signal-to-noise multiplexed detection of RNA translation throughout the whole embryo.

### Spatial profiling of RNA translation at near-genomic scale

To comprehensively assess the spatial landscape of RNA translation at near-genomic scale in early embryogenesis, we designed RIBOmap probes targeting a total of 4,096 genes (**table S1**) curated from previous bulk RIBO-seq datasets (*9*, *10*). The gene panel comprises approximately 20% of the transcriptome, with genes critical for oocytes and pre-implantation embryo development, including: (a) maternal RNAs; (b) genes involved in zygotic genome activation (ZGA); (c) genes with dynamic translation across time. We applied the optimized RIBOmap, along with simultaneous detection of cellular organelles in the same sample, to three key developmental stages in mice simultaneously: Metaphase II (MII) oocytes, late 2-cell (L2C) embryos, and 4-cell (4C) embryos (**fig. S1A**). These stages were selected to capture the major translational transitions in early development. Here, we acquired a high resolution RIBOmap dataset at a voxel size of 100 × 100 × 350 nm for a total of over 100 embryos (**Fig. 1E**) (*14*).

To achieve translatomic profiling at single-blastomere resolution in the 2-cell and 4-cell stages, we designed a 3D cell segmentation strategy that computationally segments blastomeres from intact embryo using nuclei staining as seeds, ER and mitochondria staining as embryo body masks, and cortical filament membrane staining as cell boundary masks. We confirmed by visual inspection that individual blastomeres were segmented (**fig. S1B**). Next, low-quality or partially-imaged embryos were filtered. After cell segmentation and quality control, we pooled the successfully segmented high-quality oocytes and blastomeres of all time points together for cell typing by hierarchical clustering. The RIBOmap-based translatome profile shows a clear separation of clusters by developmental time point in the UMAP space (**fig. S1C**). We found that the number of reads and genes recovered per embryo was comparable to those obtained by single-oocyte/embryo ribosome profiling (**fig. S1D-E**) (*12*), and the gene markers identified at each stage were consistent with known maternal and zygotically activated genes (**Fig. 1F and fig. S1F**). To further assess dataset reliability, we calculated the expression correlation between RIBOmap and bulk ribosome profiling data (*10*) from the same developmental stages (RIBOmap vs RIBO-Lite), benchmarking it against the correlation observed between two independent bulk datasets (RIBO-Lite vs LiRIBO-seq) (*9*, *10*). The comparable yet moderate correlation coefficients between these two comparisons (**fig. S1G**) likely reflect the intrinsic biological variability across individual embryos (*19*, *20*).

### Subcellular translatome reveals functionally diverse RNA subsets across organelles

Since localized RNA translation could drive rapid changes in local proteomes and facilitate complex assembly (*4*, *21*), we asked whether RNA translation is spatially organized within the subcellular environment of embryos and, if so, which RNA transcripts undergo such localized translation. Therefore, leveraging the organelle staining already performed in our RIBOmap experiments, we examined whether translating RNAs are preferentially enriched at specific subcellular compartments, including the nuclear periphery, mitochondria, and ER, which may serve as hubs for localized translational control, and, if the localized translation pattern changes across different developmental stages (**Fig. 2 and fig. S2**).

**Fig. 2.**
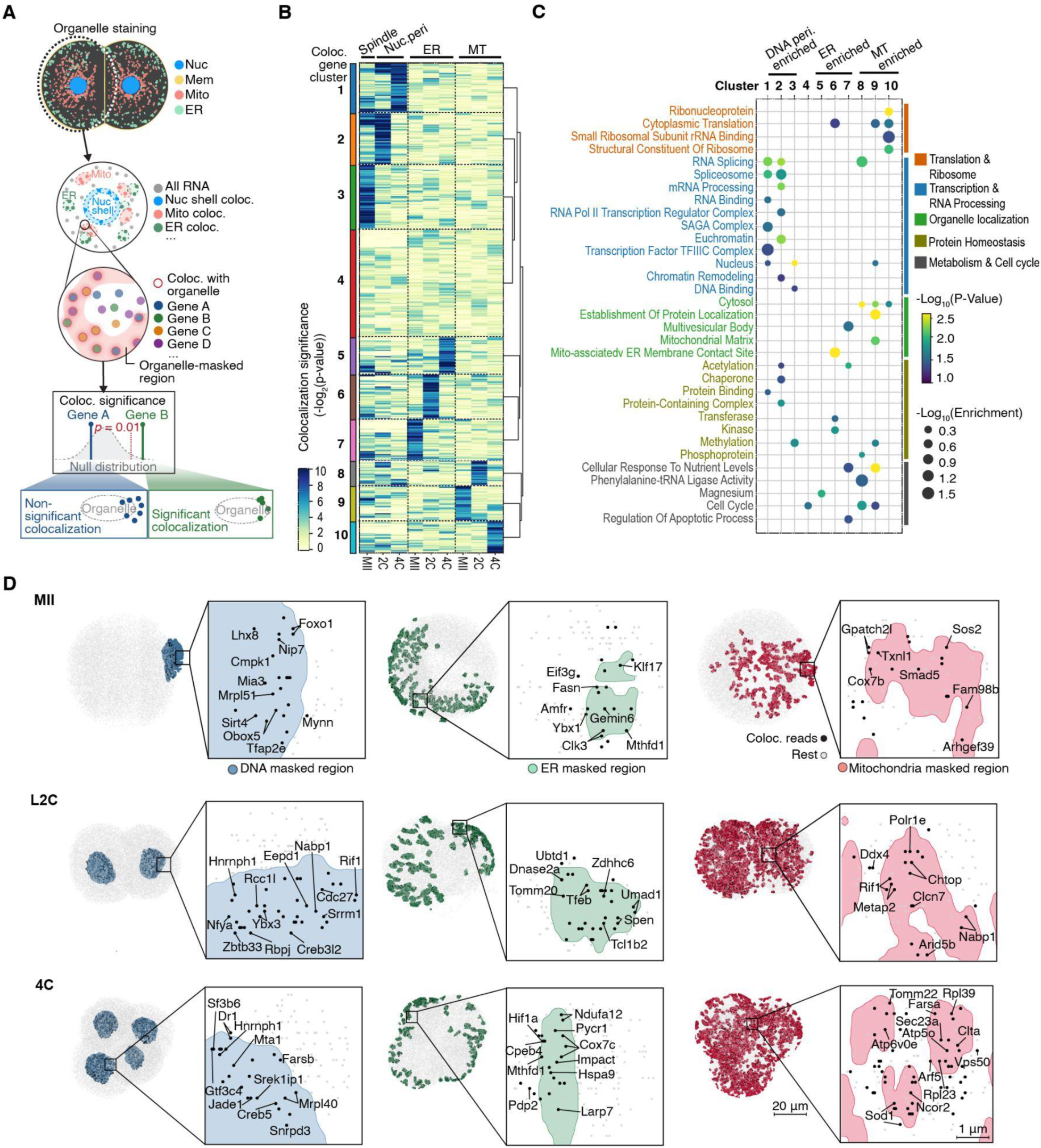
RNA-organelle colocalization reveals enrichment of functional diverse RNA subsets at nuclear periphery, ER and mitochondria. (**A**) Schematic diagram for subcellular organelle staining mask processing (for meiotic spindle or nucleus, ER and mitochondria) and RNA-organelle colocalization p-value calculation. (**B**) Heatmap showing organelle colocalization patterns across three developmental stages, with genes grouped by clustering based on their colocalization profiles, for all genes that either specifically localize to one organelle or show no significant colocalization bias. Color in the heatmap represents transformed colocalization significance values. (**C**) Dotplot showing gene ontology (GO) term enrichment of each colocalization gene cluster, with terms color-coded by broader functional category based on manual curation. (**D**) Left in each column: Global view of organelle staining for nuclear periphery (blue), ER (green), and mitochondria (red) with total RIBOmap reads in representative MII oocytes, L2C and 4C. Scale bar, 20 *μ*m. Right in each column: zoom-in views showing co-localized RIBOmap reads with selected gene names annotated. Scale bar, 1 *μ*m.

To this end, we segmented raw fluorescent organelle images from the embryos into organelle-masked regions and identified reads that optically overlapped with each mask (**Fig. 2A**). To control for intrinsic differences in gene abundance, we calculated a per-gene colocalization *p*-value at each developmental time point. This analysis revealed hundreds of genes that showed significant localization enrichment (**table S2,** colocalization *p*<0.01) for each organelle at each stage, suggesting that subcellular compartmentalization of translation is both selective and dynamically regulated. Notably, ER signals in embryos, unlike those in cultured somatic cells (*15*), were intrinsically enriched near the cortical region as distinct round-shaped aggregates (**Fig. 1E**) and showed partial overlap with mitochondrial signals, consistent with previous reports (*22*). The mitochondrial signals, on the other hand, showed a puncta-like, diffuse pattern throughout the cytoplasm, consistent with the current understanding of mitochondria morphology in embryos (*23*). We next asked whether RNAs enriched at the same organelle share coherent biological functions or instead represent functionally diverse transcript subsets. To address this, we leveraged the full matrix of colocalization *p*-values, which is comprised of three subcellular landmarks across three developmental time points for all detected genes, as input features for unbiased clustering, aiming to globally identify gene clusters with shared organelle localization patterns. The gene clustering result, based on nine colocalization values per gene (**table S2**), revealed ten colocalization gene clusters (**fig. S2A**), each exhibiting distinct subcellular distribution over time (**Fig. 2B, fig. S2B**). Specifically, Clusters 1-3 showed biased enrichment near the meiotic spindle at MII and at the nuclear periphery at the 2C and 4C stages. Clusters 5-7 showed enrichment at ER/Golgi regions, while Clusters 8-10 in mitochondrial regions. In contrast, Cluster 4 showed no specific localization bias toward any organelle or time point.

Next, we performed gene ontology (GO) analysis to systematically map the landscape of physiological pathways and biological functions associated with each identified colocalization cluster (**Fig. 2C**). Among all of the enriched GO terms, we further categorized them into five major biological categories. The breadth of these molecular and cellular functions highlights the variety and complexity of developmental processes that may also involve an unexplored layer of translational regulation at the subcellular spatial level.

Specifically, for DNA- and nuclear periphery-enriched gene clusters 1-3, genes are strongly enriched in encoding transcription regulation and RNA processing (**Fig. 2C**, **Clusters 1-3**). This finding is consistent with a previous study in cell-culture-based systems, where co-translational regulation positions splicing machinery near the nucleus for efficient nuclear import (*24*). To further examine this, we visualized the nuclear mask alongside colocalized RIBOmap reads (**Fig. 2D**), and further identified transcripts encoding transcription factors and chromatin regulators, such as *Lhx8*, *Rbpj*, *Zbtb33*, *Nfya*, *Mta1* and *Jade1*; RNA splicing and processing such as *Srrm1*, *Hnrnph1* and *Snrpd3*. Notably, genes significantly colocalized around the nucleus represent the highest proportion among all three compartments (**fig. S2C**). Together, these findings could suggest a nuclear proximity-based RNA translation that may facilitate the efficient transport of nuclear proteins back to the nucleus, a spatially-controlled RNA regulation that has not been previously reported in embryos.

Within ER/Golgi-enriched gene clusters 5-7, we observed moderate enrichment of genes associated with specialized organelle localization, including the multivesicular body (late endosome) and mitochondria-associated ER membrane (MAM) contact sites (**Fig. 2C**), potentially due to ER-mitochondria optical overlap (see discussion above). Among these, genes such as *Tfeb* and *Hspa9*, involved in lysosome biogenesis and protein quality control, respectively, were also identified (**Fig. 2D**). This suggests potential spatial and functional coupling among mitochondria, ER, and endosomal compartments. Interestingly, in contrast to somatic cells (*21*, *25*) or tissues (*15*), we did not observe strong enrichment of ER-associated transmembrane or secretory protein localization. While our gene panel includes a representative set of ER-related genes (∼10% of RIBOmap gene list), making selection bias an unlikely sole explanation, this observation may nonetheless suggest that the extracellular secretory machinery has not yet been fully established nor required at this stage of early development (*26*), though further investigation will be needed to substantiate this interpretation.

For mitochondria-enriched gene clusters 8-10, we observed enrichment of genes encoding cytosolic proteins and mitochondrial matrix components, as well as strong enrichment in ribosome biogenesis and translation, all localizing near mitochondria at the 4-cell stage (**Fig. 2C**). The enrichment in cytosolic and mitochondrial matrix proteins serves as a validation of our spatial assignments, consistent with the dispersed cytoplasmic distribution of mitochondrial staining observed across the embryo (**Fig. 1E**). Furthermore, the enrichment of translation and ribosome biogenesis machinery near mitochondria has also been shown in a previous report of mitochondria-localized translation (*21*) in cultured cells, yet has not been studied in embryos. Notably, this functional enrichment in translation and ribosome biogenesis does not become apparent until the 4-cell stage, suggesting a temporally regulated shift toward a new wave of translation that may support subsequent developmental progression.

Given the intrinsic morphological differences in organelle architecture between oocytes and embryos (**fig. S2D**), we next asked whether genes showing consistent organelle colocalization across both the 2-cell and 4-cell stages exhibit clearer organelle-specific functions. Indeed, ER-localized genes in all stages showed more defined enrichment in the Golgi apparatus and trans-Golgi network, while mitochondria-localized genes displayed stronger association with mitochondrial matrix and membrane components, both of which further validate the organelle specificity of localized translation (**fig. S2E**). In contrast, for MII oocytes, genes with cellular component annotations from current GO databases did not show comparably clear segregation into their expected organelle compartments, consistent with the fundamental differences in organelle organization between the quiescent MII state and actively dividing embryos. Taken together, we demonstrated that RNA-organelle colocalization reveals functionally diverse transcript subsets selectively enriched at the nuclear periphery, ER, and mitochondria in a stage-specific manner, with temporal shifts in localized translation representing an underappreciated layer of post-transcriptional regulation in early embryogenesis.

### Early subcellular RNA polarization in 2-and 4-cell embryos

During the first cell-fate decision in mouse preimplantation development, cells at the 8- to 16-cell stage segregate into outer polar and inner apolar populations, establishing the trophectoderm (TE) and inner cell mass (ICM) lineages, respectively. The “polarity” model (*27*, *28*) proposes that this lineage segregation arises from the differential inheritance of polarized subcellular factors during asymmetric cell divisions. Consistent with this model, blastomeres at the 8-cell stage undergo intracellular polarization, establishing asymmetric distributions of cellular components between apical and basolateral membrane domains (*29*, *30*). We therefore asked whether subsets of actively translating RNAs are already intracellularly polarized at earlier developmental stages, specifically the 4-cell and 2-cell stages, potentially priming cells for the subsequent polarization that occurs at the 8-cell stage. To address this question at a global level, we leveraged the single-molecule spatial coordinates of individual mRNAs captured by RIBOmap to characterize the subcellular spatial distribution of the translatome in intact 4-cell and 2-cell embryos.

Building on established approaches for identifying intracellular contact domains and contact-free surfaces (*31*, *32*), we first segmented each 4-cell embryo into “inner” and “outer” compartments based on the tetrahedral geometry and cell–cell contact planes (**Fig. 3A**; **video S1**), then systematically looked for RNA enrichment within each compartment. Comparative analysis of these regions identified numerous differentially expressed genes (DEGs) enriched specifically in the two segmented regions (213 genes) (**table S3**). Given the limited sample size (*N*=12 embryos), we reported nominal *p*-values to define this candidate set as Benjamini-Hochberg correction would be overly conservative in this context and risk introducing a high false-negative rate. Spatial heatmap visualization of these DEGs across multiple embryos confirms that the RNA patterns partially recapitulate the geometric segmentation, with sharp transitions aligning with the tetrahedral cell-cell contact points (**Fig. 3B**). Notably, the observed spatial patterns did not simply reflect “inner” versus “outer” distributions in an abstract sense. Rather, they resolved into two distinct subcellular patterns: (a) an embryo-edge-enriched pattern, in which transcripts are concentrated at the contact-free outer surface of the embryo, and (b) a diffuse pattern, in which transcripts are distributed broadly throughout the embryo volume. The edge-enriched pattern likely reflects active RNA anchoring or localized protection from degradation at the outer cell surface, consistent with the known asymmetry in cortical organization at this stage. The top DEGs representing each pattern are visualized in a volcano plot, where edge-enriched genes (green) display notably stronger fold changes relative to diffusely (blue) distributed genes (**Fig. 3C**).

**Fig. 3.**
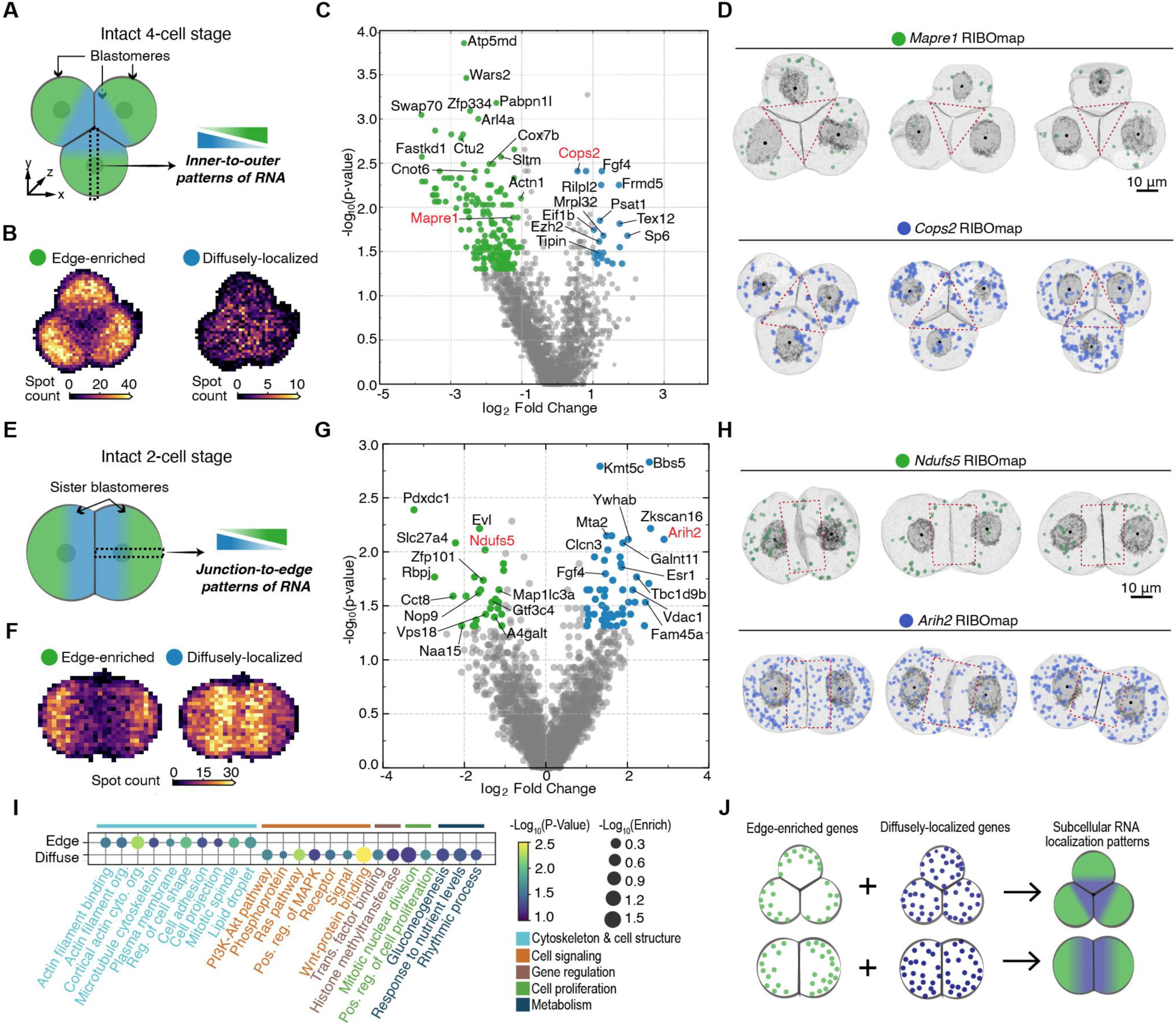
Early RNA polarization patterns in intact 2- and 4-cell embryos. (**A**) Schematic diagram of 3D inner and outer compartment segmentation in the 4-cell embryo for RNA enrichment analysis. (**B**) Aggregated spatial density visualization of genes with |log-fold| >1 in edge-enriched (left) and in diffusely-localized (right) patterns in RIBOmap across all embryos. (**C**) Volcano plots showing the differential gene expression between the two regions in 4-cell. Highlighted genes in red were selected for single-gene visualization in (**D**). Axes show (log) fold change and nominal *p*-value. (**D**) Single-gene, single-embryo visualization of RIBOmap reads for *Mapre1* (top, edge-enriched) and *Cops2* (bottom, diffusely localized), overlaid on nuclear and cell membrane staining of the same embryos in 3D from a focused-gene panel dataset. One blastomere of each embryo was removed for the purpose of visualization. Scale bar, 10 *μ*m. (**E**) Schematic diagram of 2D inner and outer compartment segmentation in the 2-cell embryo for RNA enrichment analysis. (**F**) Aggregated spatial density visualization of genes with |log-fold| >1 in edge-enriched (left) and in diffusely-localized (right) patterns in RIBOmap across all embryos. (**G**) Volcano plots showing the differential gene expression between the two regions in 2-cell. Highlighted genes in red were selected for single-gene visualization in (**H**). Axes show (log) fold change and nominal *p*-value. (**H**) Single-gene, single-embryo visualization of RIBOmap reads for *Ndufs5* (top, edge-enriched) and *Arih2* (bottom, diffusely localized) from a focused-gene panel dataset, overlaid on nuclear and cell membrane staining of the same embryos in 3D. Scale bar, 10 *μ*m. (**I**) Dotplot showing a list of GO term enrichment of differentially expressed genes in inner and outer regions shared in both 2C and 4C. Each term is colored by a broader functional category by manual curation. (**J**) Schematic diagram summarizing the two subcellular RNA patterns in 2C and 4C embryos.

Gene ontology (GO) and pathway enrichment analyses revealed that these two spatial domains are associated with distinct biological functions (**fig. S3A**). Translated RNAs enriched at the outer, contact-free surface are functionally associated with protein quality control, actin cytoskeleton regulation, and cell cycle progression. The enrichment of actin-related transcripts at the embryo periphery could be related to previous findings at the 8-cell stage, in which apical domain formation is driven by cooperative recruitment of Ezrin via the actin network to generate membrane protrusions (*30*). In contrast, transcripts localized more diffusely in the inner compartment are associated with key signaling pathways, including PI3K–Akt and Ras signaling. Prior studies demonstrated that EGFR/Ras signaling pathway prevents the mislocalization of apical-associated proteins (*33*), confirming that Ras could be active in defining early cell polarity. To validate these subcellular patterns, we performed a targeted RIBOmap experiment profiling a focused panel of eight genes to increase per-gene sensitivity, and visualized the individual gene spatial distribution in 3D alongside nuclear and membrane markers across multiple individual embryos (**Fig. 3D**). This analysis confirmed that *Mapre1*, encoding a microtubule-associated protein, is enriched at the embryo periphery, whereas *Cops2*, encoding a subunit of the COP9 signalosome, displays uniform distribution throughout the embryo (**fig. S3B**). Collectively, these results provide the first systematic evidence that subsets of actively translating mRNAs exhibit spatially polarized distributions at the 4-cell stage, potentially preceding the intracellular polarization that occurs at the 8-cell stage.

We next extended this analysis to the 2-cell stage to ask whether subcellular spatial asymmetry in RNA localization emerges even earlier. While evidence directly linking binary regionalization at the 2-cell stage to later embryo polarity remains limited, such early organization may influence cleavage orientation during division to the 4-cell stage (*34*). Rather than an inner–outer framework, we defined a junction-to-edge long axis within intact 2-cell embryos, bisecting each blastomere into an inner half (contact region near the cell junction) and an outer half (non-contact region near the cell edge) (**Fig. 3E**; **video S2**). Spatial heatmap visualization of candidate genes recapitulates the junction-to-edge segmentation, and as at the 4-cell stage, transcripts again resolve into edge-enriched and diffuse patterns (**Fig. 3F**).

Differential gene expression analysis between these segmented regions identified numerous candidate DEGs (78 genes, **Fig. 3G**) (**table S4**), demonstrating that spatial asymmetry in RNA localization emerges even earlier than previously appreciated. As with the 4-cell stage, nominal *p*-values were used to identify high-confidence spatially variable genes given the sample size constraints (*N*=13 embryos). Here, edge-enriched RNAs were associated with the cytoskeleton, regulation of cell shape, and RNA processing, whereas diffusely distributed transcripts were enriched in DNA damage response and metabolic processes (**fig. S3C**). Similar to the 4-cell findings, this suggests that the 2-cell blastomere may likewise harbor localized protein synthesis for intracellular transport, potentially supporting coordinated cell–cell communication and cleavage. We further validated this polarity by mapping representative transcripts individually, confirming *Ndufs5* displays an edge enrichment, while *Arih2* exhibits a diffuse pattern (**Fig. 3H, fig. S3D**).

Finally, we identified genes that share edge-enriched or diffuse patterns at both the 2-cell and 4-cell stages and examined their collective biological functions. Interestingly, GO analysis revealed a clearer functional distinction between the two groups: edge-localized genes are strongly and consistently enriched for actin, cytoskeleton, and cell structure-related functions, whereas diffusely distributed genes span more diverse biochemical roles in cell signaling, cell cycle regulation, and metabolism (**Fig. 3I**). Taken together, these findings indicate that RNA translation of structural, cortical, and adhesion proteins is preferentially enriched near the cell periphery across early embryonic stages, potentially establishing the molecular foundation for subsequent membrane asymmetry and cortical specification. This observation echoes a previous study in cardiomyocytes in which RNA translation was spatially restricted to cytoskeletal structures (*35*), but demonstrates that such localized translation operates as early as the cleavage-stage embryo. Moreover, the concentration of edge translated transcripts in actively dividing cells suggests a functional necessity to rapidly assemble and maintain cell boundaries and polarity during each division. This early spatial bias may therefore prime molecular asymmetries that subsequently shape blastomere behavior, intercellular coordination, and lineage specification (**Fig. 3J**).

### Harmonics reveals asymmetric subcellular patterning in MII oocytes

Our finding of two distinct subcellular RNA localization patterns in 2-cell and 4-cell embryos raises an intriguing follow-up question: does intracellular RNA biased localization exist even earlier in development? A previous proteomic study (*36*) identified intracellular spatial asymmetries emerging as early as the zygote stage, prompting us to ask whether RNA patterns might already be present at the earlier MII stage. To address this, we developed Harmonics, a graph signal processing-based framework (*17*), for unsupervised discovery of subcellular RNA patterns using RIBOmap data, without imposing preliminary assumptions about localization (**fig. S4**).

Harmonics works by decomposing spatial gene patterns into a set of underlying graph patterns, analogous to how complex sounds can be broken down into their component frequencies. The method begins by connecting spatial coordinates to create a matrix that captures the structure of connectivity (**fig. S4A**). The eigenvectors of this matrix represent wave-like patterns across the spatial graph, functioning as spatial frequencies: low frequencies capture broad, regional patterns while high frequencies capture fine-scale, localized variations, forming a basis set of spatial patterns (**fig. S4B**). For each gene, we project its transcript locations onto this frequency basis to obtain a spectrum of weights indicating how much each frequency contributes to the gene’s pattern. We then filter out high-frequency noise, retaining primarily low-frequency components, and transform back to coordinate space to yield a smoothed spatial pattern per gene (**fig. S4C**). We then use PCA and k-means clustering to extract the dominant spatial patterns. To validate this framework, we tested Harmonics on four simulated benchmark datasets (**fig. S4D**), and further applied it to a STARmap HeLa cell dataset (*15*). We successfully recovered three expected subcellular compartments: the nucleus, endoplasmic reticulum (ER), and cytoplasm from the spatial RNA distributions (**figs. S4E–G**).

Subsequently, we applied Harmonics to MII oocytes profiled by RIBOmap, and we revealed a distinct and reproducible spatial pattern across all oocytes that resembles a distinction between a diffuse interior and a peripheral “edge” at the cell surface (**Fig. 4A**). We next confirmed the spatial patterns were stable and statistically robust rather than computational artifacts (**figs. S4H-K**). In the 2D-projection spatial density visualization (**Fig. 4B**), we note that the patterns are broadly consistent with the observation profiled by Stereo-cell as “aggregation” and “dispersion” gene modules in oocytes (*37*). However, we found a 3D reconstruction revealed a more complex structure: rather than forming a complete shell, the outer “shell” region asymmetrically covers a continuous portion of the cell surface, creating a polarized distribution (**video S3**). Since the meiotic spindle of MII oocytes is positioned at the cortical edge, the 3D spatial polarization is likely organized around this asymmetrically positioned chromatin center. Notably, these binary subcellular patterns echo those observed in 2-cell and 4-cell embryos, suggesting that similar post-transcriptional mechanisms governing RNA transport or local degradation may be conserved across these early developmental stages.

**Fig. 4.**
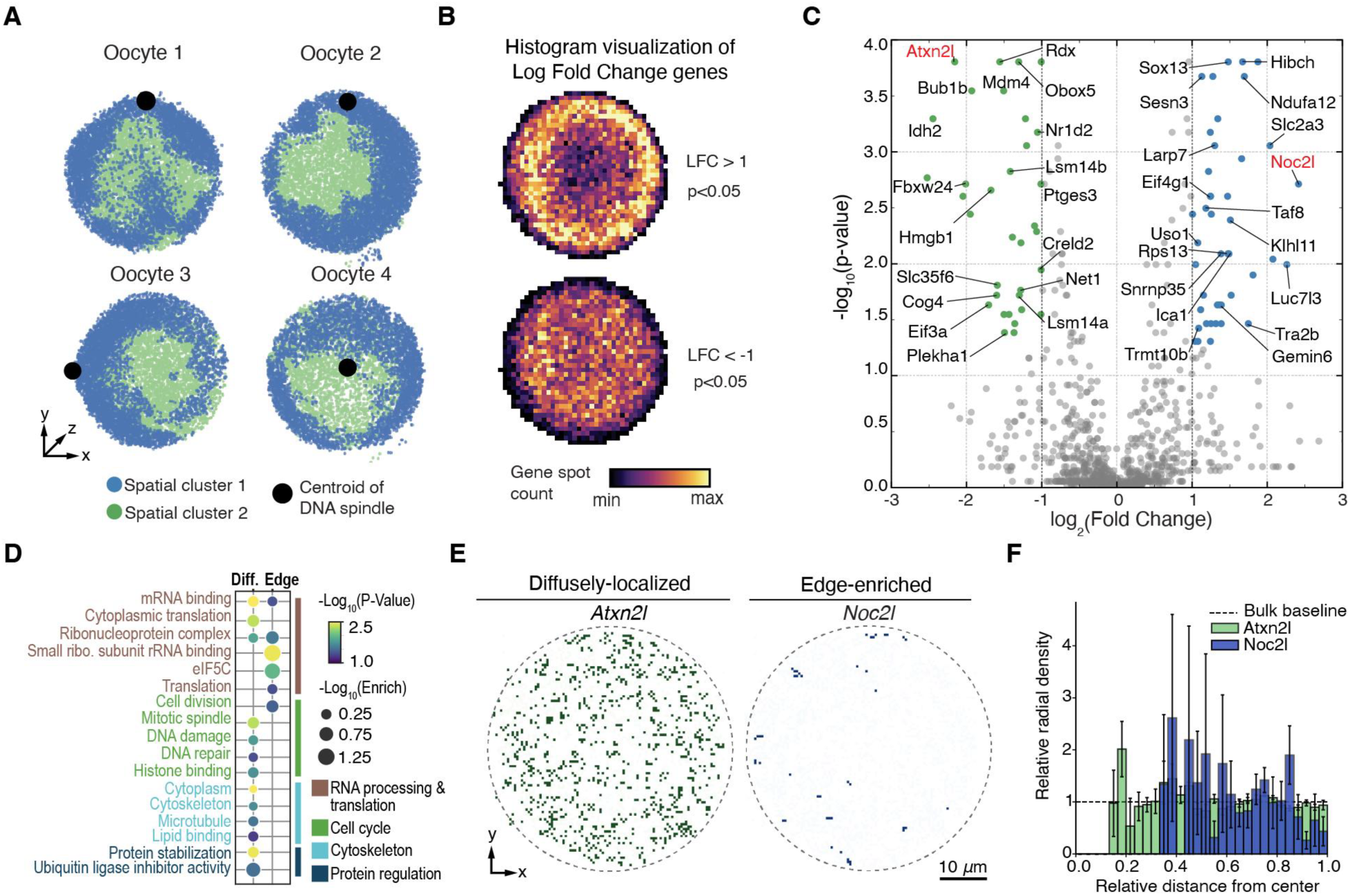
3D spatial asymmetry in MII oocytes identified by Harmonics. (**A**) Spatial assignment of two Harmonics clusters in oocyte space and can be visually detected as “diffuse interior” and off-center “shell” patterns. Four representative oocytes are shown *(N*=15-20). Green and blue represent two Harmonics clusters, respectively. (**B**) Aggregated spatial density visualization of genes with |log-fold| >1 and *P*<0.05 in each subcellular compartment. (**C**) Volcano plot showing the differential gene expression of RIBOmap enriched in each of the two regions. Top genes are highlighted in color and annotated. (**D**) Dotplot showing a list of GO term enrichment of differentially expressed genes in edge and diffuse patterns in MII oocytes. Each term is colored by a broader functional category by manual curation. (**E**) RIBOmap 2D-projected visualization of one diffuse gene (*Atxn2l*) and one edge-enriched gene (*Noc2l*). Scale bar: 10 *μ*m. (**F**) The relative radial intensity of normalized *Atxn2l* and *Noc2l* RIBOmap signals against eight other genes (bulk) within each radial bin from quantification of the radial distribution.

We identified hundreds of DEGs with significant association in each region (**Fig. 4C**, **table S5**). The edge-enriched genes are moderately associated with functions in RNA translation machinery. In contrast, the interior-enriched genes are predominantly associated with cytoskeleton and cell structures, cell cycle, translation and protein regulation, therefore spanning a wider range of functions from maintaining cell structure to basic cell metabolism (**Fig. 4D**). To validate the diffuse vs edge patterns, we selected three representative genes (*Atxn2l*, *Idh2*, and *Noc2l*) and repeated a small-scale RIBOmap experiment, which confirmed their respective spatial distributions (**Figs. 4E-F**).

Intriguingly, cytoskeleton-related RNAs, such as *Atxn2l*, display a diffuse cytoplasmic distribution in MII oocytes, suggesting that transcripts essential for cellular architecture are broadly dispersed throughout the oocyte at this stage. This is consistent with the known biology of MII oocytes, in which maternal mRNAs are largely held in a translationally dormant state and stored in cytoplasmic structures including cytoplasmic lattices (*38*, *39*), serving as reservoirs for later developmental needs. This stands in stark contrast to the edge-enriched localization of functionally similar genes observed in 2-cell and 4-cell embryos (**Fig. 3I**), which may point to a spatiotemporal shift in the subcellular distribution of cytoskeletal RNA translation following ZGA, from diffuse cytoplasmic storage in the meiotically arrested oocyte to peripherally concentrated translation once active cell division starts and the timely establishment of cell boundaries and polarity becomes critical. Therefore, this spatiotemporal reorganization of cytoskeletal RNA translation between these stages and whether this change is triggered by fertilization or ZGA represents an interesting open-ended question requiring future mechanistic investigation.

### Harmonics reveals distinct translational asymmetries in 2-cell and 4-cell embryos

Our molecular understanding of 2-cell asymmetry in mouse embryogenesis has evolved significantly from the traditional “equivalent hypothesis” (*40*, *41*), which assumes blastomeres are identical, toward an “asymmetric hypothesis” that recognizes their differences. Accumulating evidence suggests inter-blastomere differences across multiple molecular layers, including epigenetic regulation (*42–44*), transcriptome (*45*) and proteome (*36*), that bias future fate toward either ICM or TE. However, blastomere asymmetry has not been systematically analyzed at the level of mRNA translation, which may be critical for understanding and even predicting the first cell fate decision. Moreover, imaging-based molecular profiling offers a robust technical approach for identifying key gene differences without dissociating embryos, enabling us to trace individual blastomeres to their embryo of origin without unnecessary experimental complications.

A foundational challenge in this analysis was developing a consistent method for assigning A/B blastomere identity across all embryos, since previous studies have noted that reproducible inter-blastomere differences are difficult to distinguish from the substantial inter-embryo variation present at these early stages (*19*, *20*). Additionally, we were motivated to explore whether we could directly connect asymmetric differences in 2-cell to the more enhanced cell-to-cell differences that emerge at the 4-cell stage (*32*, *36*, *45*), thereby tracing how early translational asymmetry evolves as development proceeds.

Building on the unsupervised nature of the Harmonics framework (**figs. S4A-D**), we extended it to investigate spatial pattern differences across multi-cellular space by applying it jointly to the RIBOmap dataset of 2-cell and 4-cell embryos. Interestingly, at k=2 clustering, Harmonics produced well-defined separations: two distinct blastomeres in 2-cell embryos and two pairs of blastomeres in 4-cell embryos, and the patterns are linked between the two stages (**videos S4-5**). To validate these groupings, we confirmed the robustness against slight parameter perturbations and background shuffling controls (**figs. S5A–C**). Given that consistent inter-blastomere differences could be detected across embryos, we used clusters 0 and 1 (**fig. S5A**) to designate 2-cell blastomeres as sister blastomeres A and B (**Fig. 5A**), and in the two 4-cell pairs as pair A and pair B (**Fig. 5D**), respectively. Together, these results suggest that a subtle translational asymmetry already present at the 2-cell stage can be connected to distinct sister-blastomere pairs at the 4-cell stage. By applying the same analytical framework across both developmental timepoints, asymmetry patterns in RNA translation can be directly compared to reveal how early molecular differences evolve as development proceeds.

**Fig. 5.**
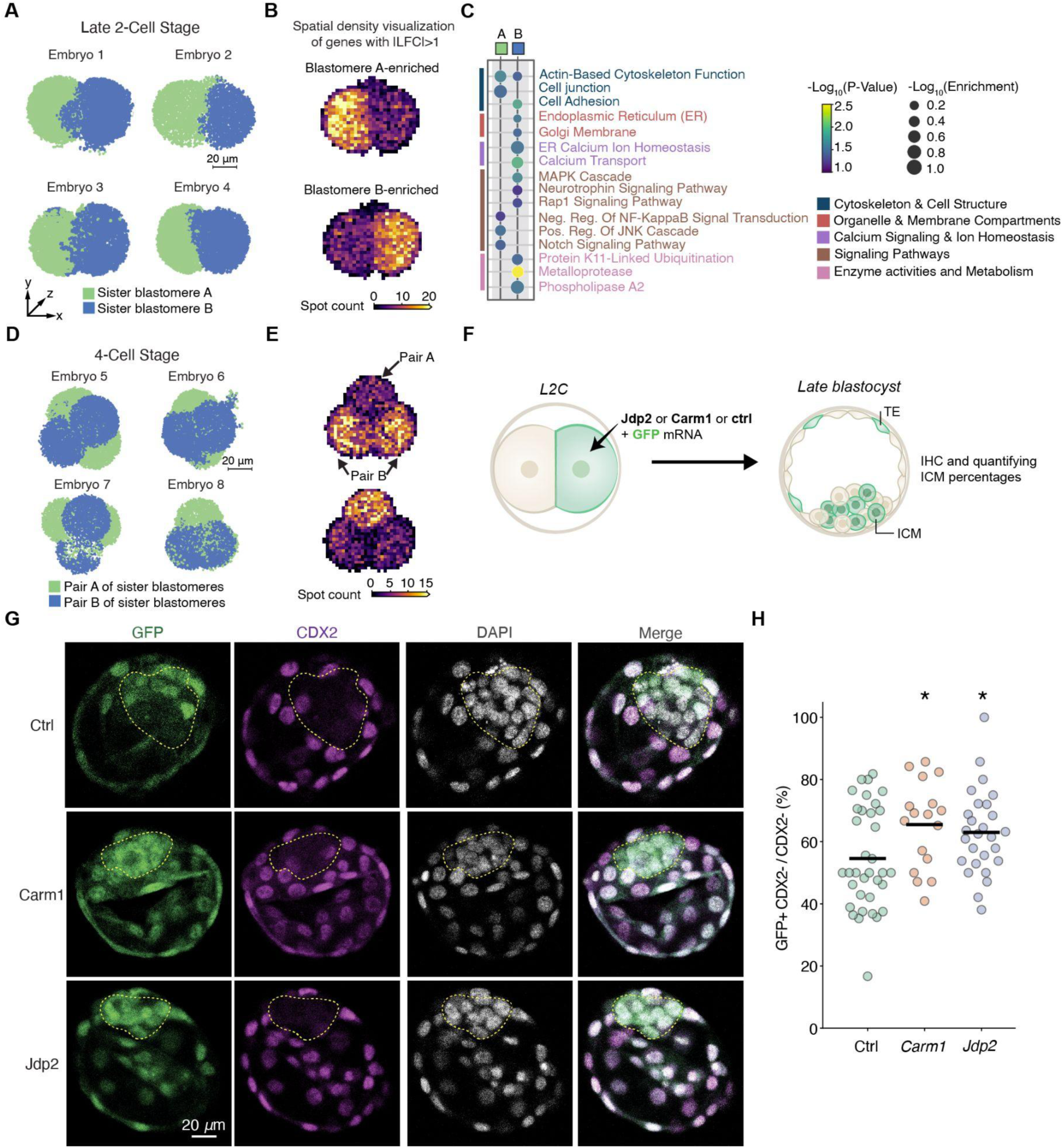
Unbiased pattern discovery identifies stage-specific translational asymmetries in 2- cell and 4-cell embryos. (**A**) Spatial assignment of two Harmonics clusters in late 2-cell embryos (n=8 embryos; four representative embryos are shown). Green and blue represent the two clusters, corresponding to blastomere A and B, respectively. Clusters were jointly defined using both late 2-cell and 4-cell samples (**A** and **D**), enabling consistent color assignment across stages. (**B**) Aggregated spatial density visualization of genes assigned to each cluster in (**A**) with |log-fold| >1 in 2-cell samples. (**C**) Dotplot showing a list of GO term enrichment of differentially expressed genes in asymmetric blastomeres of 2-cell. Each term is colored by a broader functional category by manual curation. (**D**) Spatial assignment of two Harmonics clusters in 4-cell embryos (n=6 embryos; four representative embryos are shown). Green and blue represent the two clusters representing two distinct pairs of blastomeres and corresponding to blastomere pair A and B, respectively, and are directly comparable to the cluster assignments in (**A**). (**E**) Aggregated spatial density visualization of genes assigned to each cluster in (**D**) with |log-fold| >0.5 in 4-cell samples. (**F**) Schematic of the overexpression strategy. One blastomere of 2-cell embryos was injected with control RNA or mRNA encoding GFP-tagged *Carm1* (positive control) or *Jdp2*. GFP serves as a lineage tracing marker to track the progeny of the injected blastomere. Embryos were cultured to the late blastocyst stage, and the contribution of injected cells to the ICM lineage was assessed by quantifying the proportion of GFP-positive and CDX2-negative cells by immunostaining. (**G**) Representative immunostaining images of control, Carm1-OE, or Jdp2-OE blastocysts, showing GFP, CDX2 and DAPI channels. Scale bar, 20 *μ*m. (**H**) Dot plot showing the CDX-negative and GFP-positive proportions of total GFP-positive cells. Each dot represents an embryo. Both Carm1-OE and Jdp2-OE show increased ICM contribution compared to the control. Student t-test, \**P*<0.05.

Having assigned A/B blastomere identities at the 2-cell stage and A/B pair identities at the 4-cell stage using Harmonics, we next identified differentially translated genes between A and B groups (absolute log_2_-fold change > 1) at both stages (**table S6**, **figs. S6A–B**), and visualized their distributions using 2D projection spatial density plots (**Figs. 5B, E**). Comparing the differentially translated genes across stages, we found that approximately 25% of genes in either the A or B group are shared between the two timepoints (**fig. S6C**), suggesting partial conservation of translational asymmetry in development. Focusing on the functional significance of 2-cell asymmetry, GO analysis revealed that blastomere A was specifically enriched in signaling pathways including NF-κB, JNK, and Notch, whereas blastomere B showed broader enrichment across diverse biochemical processes such as ER/Golgi function, calcium transport, protein degradation, and enzyme metabolism (**Fig. 5C**). Notably, these findings align with a recent proteomic study identifying 2-cell asymmetry in protein degradation and ER function (*36*), thereby connecting our observations of translational asymmetry to downstream protein-level differences.

Among the asymmetrically translated genes, *Jdp2* stood out as being consistently enriched in the B group at both the 2-cell and 4-cell stages (**figs. S6A–B**), which prompted us to investigate its functional role in early development. To validate our asymmetry analysis more broadly, we designed a focused RIBOmap panel of eight genes, including *Jdp2* and *Carm1*, the latter has previously been shown to exhibit asymmetric function as early as the 2-cell stage (*42*). Using this smaller gene panel, blastomeres from seven embryos formed two well-separated clusters in UMAP space, with the two blastomeres of each embryo consistently falling into opposite clusters (**figs. S6D–E**). We confirmed that no significant differences in total read counts existed between clusters (**fig. S6F**), and that *Carm1* showed no significant expression differences between groups (**fig. S6G**), consistent with prior report (*46*). In contrast, Jdp2 displayed significant translational differences between the two clusters at the 2-cell stage (**fig. S6H**). Jdp2 is a transcription factor with established roles in cell differentiation, stress response, and proliferation (*47*, *48*), and has previously been shown to partially substitute for Oct4 in iPSC reprogramming (*49*). To test whether its translational asymmetry at the 2-cell stage carries functional consequences for cell fate, we overexpressed Jdp2 by mRNA injection into one blastomere at the 2-cell stage (**Fig. 5F**), and found that the injected cells showed increased contribution to the ICM at the late blastocyst stage similar to that of *Carm1* mRNA injection (**Figs. 5G-H**). Together, these results demonstrate the power of RIBOmap to uncover asymmetric translation in early embryos that are also functionally consequential for lineage determination.

### Translational heterogeneity in 4-cell embryos

Next, we explored whether additional heterogeneity patterns beyond the 2:2 distribution might exist at the 4-cell stage. While the 2-region (2:2) pattern emerged when Harmonics was applied at k=2, we asked whether increasing the clustering resolution to k=3 and k=4 might reveal further complexity (**fig. S7A**, **video S6**). At k=4, we identified a second reproducible and statistically significant pattern: a 3-region, or 1:1:2, arrangement (**Fig. 6A, fig. S7A**). Comparing the 2-region and 3-region assignments, we found that the 3-region pattern largely arises from a splitting of pair A into two distinct cells, which we designated clusters A′ and A′′, while the remaining two-cell group, designated cluster B′, showed substantial overlap with pair B from the 2-region assignment (**Fig. 6B**). Although the 3-region pattern achieved a lower stability score (0.75) than the 2-region pattern (1.00) in the 4K-gene RIBOmap dataset, both were statistically significant (**figs. S6C, S7A**). Consistent with this, differentially expressed genes could also be identified under the 3-region framework (**Fig. 6C**, **table S7**), supporting such asymmetric splitting. Notably, when the same eight-gene panel used to validate 2-cell asymmetry (**figs. S6D–E**) was applied to 4-cell embryos, it predominantly resulted in a 3-region, 1:1:2 clustering pattern (**figs. S7B–E**). Importantly, analysis of the *Carm1* expression revealed a 3-region pattern (**fig. S7F**), consistent with prior reports (*43,50*). These findings collectively suggest that the 4-cell stage exhibits multiple layers of heterogeneity, the precise arrangement of which may depend on the subset of genes examined.

**Fig. 6.**
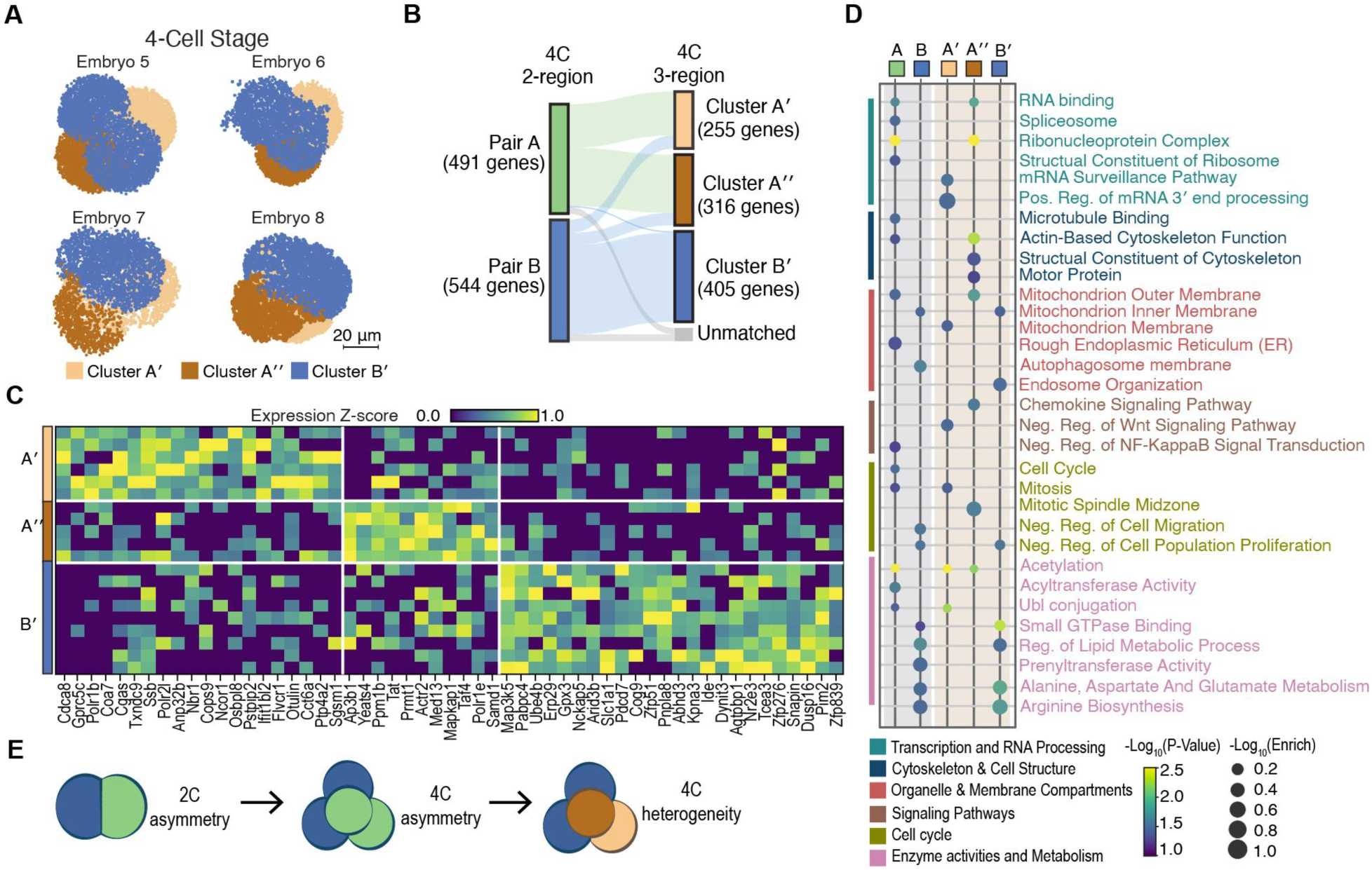
Unbiased pattern discovery identifies translational heterogeneity in 4-cell embryos. (**A**) Spatial assignment of three Harmonics clusters in 4-cell embryos (n=6 embryos; four embryos are the same as in Fig. 5D). Yellow (cluster A′), brown (cluster A′′) and blue (cluster B) represent the three assigned clusters, representing one cell each (yellow and brown) and a pair of cells (blue). (**B**) Riverplot showing the connection between 2-cluster (Fig. 5D) and 3-cluster (**A**) assignments in 4C by DEG comparison. (**C**) Heatmap showing normalized gene expression Z-scores for the most variably expressed genes across the three region assignments (clusters A′, A′′, and B′) of 4-cell embryos. (**D**) Dotplot showing a concatenated list of GO term enrichment of differentially expressed genes in asymmetric pairs from 2-region and cells from 3-region assignments. Each term is colored by a broader functional category by manual curation. (**E**) Schematic view of proposed embryo asymmetry developmental model from 2-cell to 4-cell.

To further characterize the functional significance of these heterogeneity patterns, we visualized GO analysis on both the 2-region and 3-region cell clusters at the 4-cell stage (**Fig. 6D**). In the 2:2 pattern, pair A was highly enriched for RNA splicing and translation, a signature that persists specifically in cluster A′′ of the 3-region assignment. Cluster A′′ was additionally distinguished by enrichment in cell-structure-related functions not observed in the other clusters. In contrast, pair B and its corresponding cluster B′ were characterized by enrichment in basic enzymatic activities and metabolic processes, including amino acid synthesis and metabolism (**Fig. 6D**). Together, these patterns suggest a functional division among 4-cell blastomeres that may be tied to distinct developmental trajectories or cell-cycle effect (**Fig. 6E**). Further investigation into how these differentially regulated translational programs contribute to lineage specification will be important for establishing their causal roles in early cell fate determination.

In summary, we developed and utilized a new platform of RIBOmap to systematically analyze embryonic RNA translation across three interconnected spatial scales. At the molecular-to-organelle scale (**Fig. 2**), we found that distinct subset of transcripts cluster near the nuclear surface, ER/Golgi network, and mitochondrial puncta, and that these organelle associations shift between MII, 2-cell, and 4-cell stages in a manner coupled to the biological roles of the encoded proteins. Zooming out to the subcellular-to-whole-cell scale (**Figs. 3-4**), we adopted Harmonics, a graph-signal-processing framework, and discovered a polarized interior-versus-surface partition of translating mRNAs in MII oocytes and recurrent edge-biased translation of cytoskeletal and cortical proteins in actively dividing embryos. Further out to the intercellular scale (**Fig. 5**), the same framework exposed reproducible translational asymmetry between sister blastomeres at the 2-cell stage that extend to further asymmetry as pairs in 4-cell blastomeres, with Jdp2 emerging as an asymmetrically translated RNA whose elevated expression increased the chance of the derived cell progeny toward ICM. Lastly, we closely examined the translational heterogeneity in 4-cell blastomeres and revealed a more complex patterning at 4-cell stage (**Fig. 6**). Together, these multiple layers of spatial regulation constitute an integrated, three-dimensional view of how ribosome-engaged RNAs are organized from organelle contacts, through cell-wide geometry, to blastomere-to-blastomere asymmetry.

## Acknowledgments

We thank Kamal Maher for providing the initial computational framework for Harmonics, Daoyuan Qian for a helpful comment on parameter tuning, Jiahao Huang and Zefang Tang for assistance in RIBOmap data processing. We thank Jiakun Tian for assistance in RIBOmap probe design. We also thank Mengyao Li, Jiakun Tian, Jiunn Song and Qianying Yang for thoughtful comments on the manuscript.

## Funding

National Institutes of Health grant R01HD116750 (YZ)

National Institutes of Health grant DP2 1DP2GM146245-01 (XW)

Howard Hughes Medical Institute (YZ)

Edward Scolnick Professorship (XW)

Merkin Institute Fellowship (XW)

Packard Fellowship (XW)

Sloan Research Fellowship (XW)

Helen Hay Whitney fellowship of HHMI (JR)

## Author contributions

Conceptualization: YZ, XW

Methodology: XW, JR, SF, HZ

Investigation: JR, SF, CZ, YH, HZ

Visualization: JR, SF, HZ, YH

Funding acquisition: YZ, XW

Project administration: YZ, XW

Supervision: YZ, XW

Writing – original draft: JR, SF

Writing – review & editing: JR, SF, XW, YZ, CZ, YH, HZ

## Data, code, and materials availability

RIBOmap datasets of one 4,096-gene and two 8-gene in embryos in Zenodo (https://zenodo.org/records/19544184?preview=1&token=eyJhbGciOiJIUzUxMiJ9.eyJpZCI6IjN jMWVjMGY3LWY4YzYtNGY2My1hYTQ0LWRjMmQyMjI4ZWE4NSIsImRhdGEiOnt9LCJyYW5kb20iOiI4NzUzNDg2MzdlOTZkM2YwODEwMTE3NjU3MDBiM2UxMyJ9.8HzIHzrMr ZgUqkFJen4Fk8vwRDNbCJFCTaxy2WiHWetaVIirnFtH5L3ncqyNKBzVtGgJDljFSKBSJgJQn eZZSQ).

The specific gene lists, as well as their organelle-colocalization *P*-values, polarity and asymmetry DEG parameters are provided as supplementary tables (**tablesx S2-7**). Two bulk RIBO-seq expression data for the same developmental stages were accessed from C. Zhang et al (DOI: 10.1126/sciadv.abj3967) and Z. Xiong et al (DOI: https://doi.org/10.1038/s41556-022-00928-6).

This work was implemented based on MATLAB v.R2020b, Python v.3.9.23 and R v.3.6.3. The following packages and software were used in data analysis: ImageJ v.1.51, anndata v.0.7.5, PyGSP v0.6.1, matplotlib v.3.1.3, seaborn v.0.13.2, scanpy v.1.10.1, numpy v.1.19.4, scipy v.1.13.1, pandas v.1.3.5, scikit-learn v.1.6.1, numba v.0.54.1, tifffile v.2021.7.2, scikit-image v.0.18.3, Seurat v.3.2.257, SeuratDisk v.0.0.0.9013, ggplot2 v.3.3.5, factoextra v.1.0.7, dplyr v.1.0.4, circlize v.0.4.13, IRanges v.2.20 and Picky 2.2.

## Supplementary Materials

Materials and Methods

Figs. S1 to S7

References (*14–18, 52–55*)

Tables S1 to S7

Movies S1 to S6

